# Genome assembly of five Tephritid species for the enhancement of the Sterile Insect Technique

**DOI:** 10.1101/2025.09.15.670340

**Authors:** Haig Djambazian, Shu-Huang Chen, Pierre Bérubé, Katerina Nikolouli, Maria-Eleni Grigoriou, Dimitris Rallis, Alistair Darby, Nikolai Windbichler, Marc F. Schetelig, Philippos Aris Papathanos, Kostas Mathiopoulos, Gur Pines, Jiannis Ragoussis, Kostas Bourtzis

## Abstract

Tephritidae insect pests account for extensive crop damage and yield losses globally. Modern, sustainable pest management approaches are species-specific and, therefore, high-quality genome assemblies are required for their application. Here, we present chromosome-level assemblies for five members of the Tephritidae family: *Anastrepha fraterculus, Anastrepha ludens, Bactrocera dorsalis, Bactrocera zonata* and *Zeugodacus cucurbitae*. The assemblies used long read sequencing polished with short read sequencing and scaffolded using Hi-C (chromatin conformation capture) sequencing. Prior to scaffolding the assembly deduplication was performed to separate a primary assembly and an alternate assembly, and each was then scaffolded independently. The scaffolded assemblies reached N50 length in the range of 60Mb to 120Mb. The scaffolded assemblies were verified with BUSCO and completeness was in the range 97% to 98.5% and had very low duplicated, fragmented and missing orthologs.

## Introduction

Members of the Tephritidae family that cause significant yield and economic losses are worldwide distributed in areas that include: tropical areas of Southeast Asia and in the sub-Saharan region in Africa (*Bactrocera dorsalis*), the Southern and South-East Asian countries and the Arabian Peninsula (*Bactrocera zonata*), Asian and Australian-Oceanian Regions (Z*eugodacus cucurbitae*), Central America (*Anastrepha ludens* and *Anastrepha fraterculus*) and in the Southern United States, Caribbean Islands, and South America (*Anastrepha fraterculus*) (EPPO 2024). The spread of these species can occur either through natural movement of the flies or by human-assisted means of transport (imports of fruit commodities or fruit in passenger luggage) thus elevating the risk of invasion in new areas. Additionally, the ongoing climate crisis is expected to provoke the spread of invasive insects into new regions (Papadopoulos et al. 2024; Qin et al. 2019; Ullah et al. 2023).

*Bactrocera dorsalis* (Hendel), the oriental fruit fly, belongs to the “dorsalis complex” which includes pest species that cause devastating fruit losses in global fruit production. Due to its extensive host range (more than 270 plant species) (Vargas et al. 2015), its high reproductive rate and the continuously rising global temperatures, it is estimated that previously unsuitable areas are or will soon be available for the introduction and establishment of *B. dorsalis* (Jaffar et al. 2023; Zhao et al. 2024).

*Bactrocera zonata* (Saunders), the peach fruit fly, mainly attacks peach and guava cultivars but mango and citrus fruits are also among its long list of cultivated hosts (Delrio & Cocco 2012; White & Elsonharris 1994). *Bactrocera zonata*’s polyphagous habits and the absence of dormancy allow the pest to be continuously active and complete several generations throughout the year (El-Mahdy et al. 2009; Khan & Naveed 2017).

*Zeugodacus cucurbitae* (Coquillett), the melon fruit fly, is considered native to India but it has been introduced in several Asian and African countries, as well as in regions of Oceania (De Meyer et al. 2015). Cucurbit crops are the main hosts of *Z. cucurbitae*, although non-cucurbit crops have also been reported (Dhillon et al. 2005). *Zeugodacus cucurbitae* has been shown to withstand a wide range of temperatures at all developmental stages, which can enhance its geographical distribution potential (Ahn et al. 2022).

The Mexican fruit fly, *Anastrepha ludens* (Loew), is a major pest of citrus and mango in tropical and subtropical areas (CABI 2019). It is considered one of the most abundant fruit flies in these areas with substantial economic impacts that range from fruit losses to increased production costs and trade restrictions (Hernández et al. 2017; Zapata 2022).

Another member of the *Anastrepha spp*., the South American fruit fly, *Anastrepha fraterculus* (Wiedemann), is a highly destructive pest of a wide range of fruits including citrus, apples and stone fruits (Ovruski et al. 2003; Segura et al. 2006). Due to its extent morphological and genetic variability, *A. fraterculus* is considered a cryptic species complex, a factor that has contributed to enhanced quarantine regulations and international trade regulations (Hendrichs et al. 2015).

The need to develop efficient pest management techniques against the above fruit flies along with the global shift away from chemical applications have advanced the sterile insect technique (SIT) into a prime approach. The SIT is based on the release of irradiated sterile males in the wild that mate with wild females resulting in a decrease of the population in the field (Dyck et al. 2021). It is an environmentally friendly and sustainable control method that has been applied for the suppression, eradication, prevention or containment of several insect pests that affect economically important crops, livestock, and human health (Dyck et al. 2021; Klassen et al. 2021).

SIT application efficiency and cost-effectiveness is improved when genetic sexing strains (GSS) of the target pest are employed. Male-only releases have been previously proven to increase the SIT effectiveness and decrease the overall cost of the technique (Hendrichs et al. 1995; Rendón et al. 2004). Genetic sexing strains of fruit flies like *Ceratitis capitata* have been developed through irradiation and classical genetic approaches and are used worldwide with a proven success record (Franz et al. 2021; Augustinos et al. 2017). These GSSs are based not only on visible markers (white pupae - wp phenotype), but also on conditionally lethal traits (temperature-sensitive lethal - tsl phenotype), which allow for the removal of females early in the developmental process (Franz et al. 2021; Caceres 2002). The identification of selectable markers for the current GSS, like VIENNA 7 and VIENNA 8 of *Ceratitis capitata*, was achieved by classical genetic approaches and lasted over two decades (Franz et al. 2021; Augustinos et al. 2017). The continuous introduction and establishment of insect pests (Zingore et al. 2020) have urged scientific efforts to find faster and transferable approaches for the identification of new selectable markers across different species (FAO/IAEA 2019, 2021; Ward et al. 2021; Sollazzo et al. 2024).

Recent efforts have been focused on a generic (neo-classical) approach that is based on the construction of non-transgenic GSS for SIT applications (FAO/IAEA 2019, 2021). The generic approach involves the identification of genes and their specific mutations responsible for desirable traits in a species that can function as selectable markers. Induction of mutations in the orthologous genes of other species and the linkage of the wild-type allele of the gene to the male sex can lead to the construction of a new GSS for an SIT target species (Ward et al. 2021; Sollazzo et al. 2024).

A high-quality genome sequencing and assembly can facilitate the identification of genes and their unique polymorphisms and advance the application of the generic approach in several species. This will subsequently allow for efficient genome editing and prevention of off-target changes through the implementation of genome editing approaches, such as the CRISPR/Cas9 technology (Bai et al. 2019; Komal et al. 2023; Paulo et al. 2022; Sollazzo et al. 2024; Ward et al. 2021). The CRISPR/Cas9 technology targets predetermined locations in the genome and therefore genomic data of high accuracy are required for efficient editing and mutation induction. In addition, a highly precise Y-chromosome assembly of the species used for GSS development can aid in the identification of suitable regions for knock-ins that will link the wild type allele of the selectable marker to males (FAO/IAEA 2019, 2021).

In this study we sequenced and assembled at the contig level, male samples of five Tephritid fruit flies. Using state-of-the-art genome sequencing technologies (Illumina NovaSeq; Oxford Nanopore and PromethION) a valuable resource of genomic data with improved quality and accuracy was created. These new genome assemblies can be used for the identification of selectable markers and the induction of mutations in these five fruit fly species, while the assembly of the Y-chromosome further supports the development of GSS. Since the samples used for sequencing were in all cases males, valuable data for the Y-chromosome are also offered that will assist in the identification of suitable locations for gene insertions in this chromosome. Although all our assemblies are at the contig rather than the chromosome level, they bear value for population structure and global invasion route studies and offer essential data for studying the evolutionary history of the species (Zhang et al. 2023). These contig level assemblies will serve as an index for pinpointing the best targets for genetic manipulation and development of tephritid fruit flies GSS.

## Results

The assembly workflow produced high quality contig-level genomes using PacBio (continuous long reads) and Nanopore long reads based on BUSCO results. The Hi-C sequencing produced chromosome-scale scaffolds for all the genomes (See Table 1 and Figure 1). The assembly workflow also yielded highly complete haploid principal assemblies. The long read polished assemblies have a >97% complete BUSCO detected (Table 2). The duplicate purging process also successfully removed duplicated sequences present in the initial long read assembly. The Hi-C maps shown in Figure 2 show a very clean picture of the scaffolding with no apparent mis-assemblies. The repeat annotation (Table 3) shows the repeat types and numbers detected in the assemblies. The number of base repeats masked in each of the genomes ranges from 3.17% to 3.7%.

**Table 1.** Genome statistics for different, final principal (haploid) assemblies are shown. The scaffolded assemblies all reach an N50 length ranging from 62 Mb to 121 Mb with three to four scaffolds, which is indicative of a successful scaffolding process. Only *Zeugodacus cucurbitae* scaffolded to a smaller than expected genome size.

| Statistics | *Anastrepha fraterculus* | *Anastrepha ludens* | *Bactrocera dorsalis* | *Bactrocera zonata* | *Zeugodacus cucurbitae* |
| --- | --- | --- | --- | --- | --- |
| # contigs | 4,973 | 1,238 | 588 | 660 | 5,140 |
| # contigs (>= 50000 bp) | 67 | 403 | 58 | 162 | 239 |
| Largest scaffold | 182,068,020 | 125,331,998 | 102,926,020 | 103,247,910 | 70,658,899 |
| Total length | 793,193,562 | 720,811,084 | 537,310,284 | 600,442,075 | 374,778,270 |
| Total length (>= 50000 bp) | 766,717,392 | 706,688,108 | 531,140,571 | 593,885,298 | 349,303,203 |
| N50 | 121,046,886 | 118,249,840 | 76,567,395 | 65,592,266 | 62,275,020 |
| L50 | 3 | 3 | 4 | 4 | 3 |
| GC (%) | 37 | 37 | 37 | 36 | 35 |
| # N's | 225,200 | 231,400 | 105,800 | 118,600 | 134,300 |
| NG50 | 109,980,515 | 114,208,257 | 40,758,100 | 45,696,414 | - |
| LG50 | 4 | 5 | 8 | 7 | - |

**Table 2.**
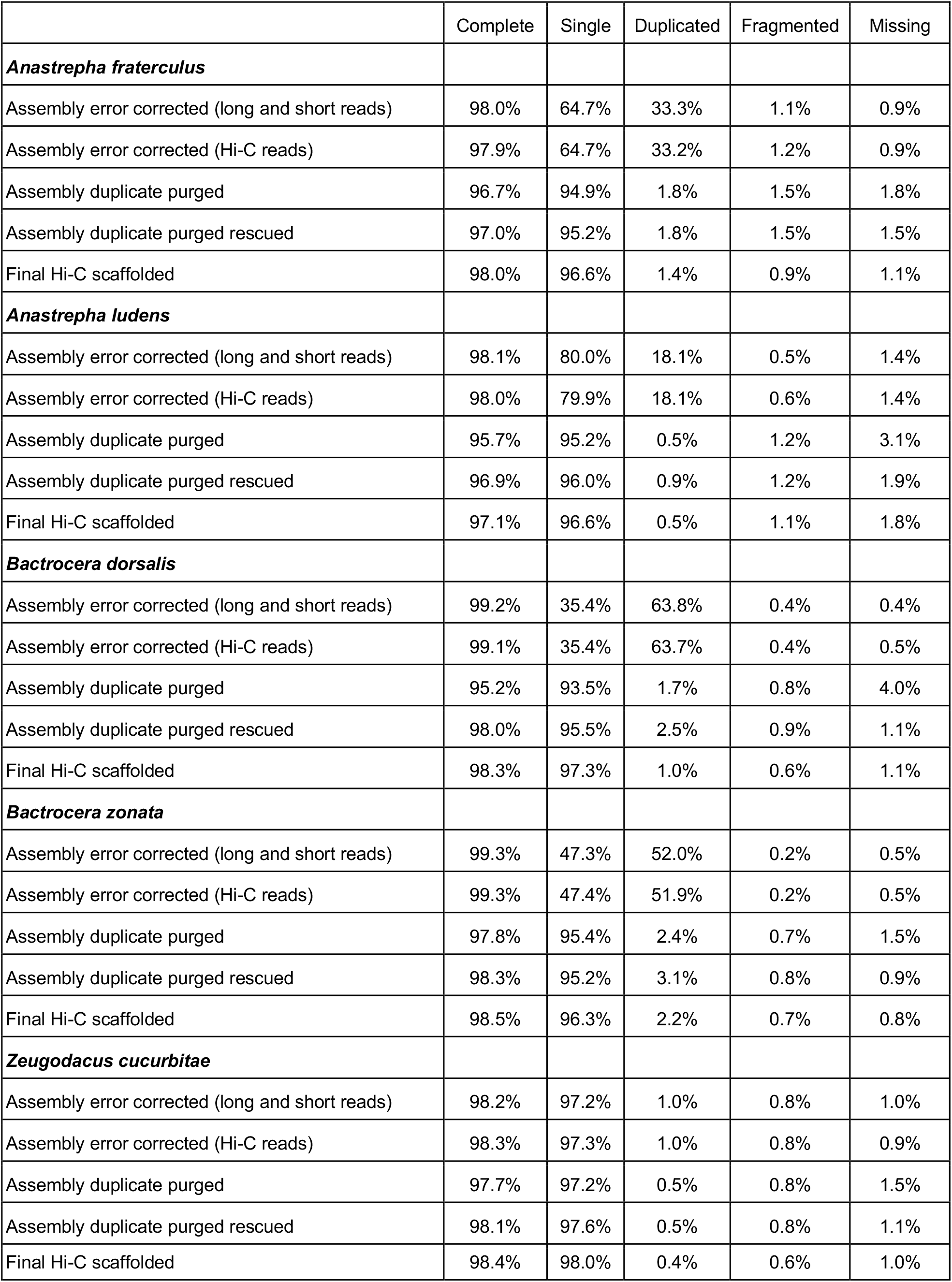
BUSCO statistics for the principal assemblies at the different stages of the assembly process (using diptera_odb10).

|  | Complete | Single | Duplicated | Fragmented | Missing |
| --- | --- | --- | --- | --- | --- |
| ***Anastrepha fraterculus*** |  |  |  |  |  |
| Assembly error corrected (long and short reads) | 98.0% | 64.7% | 33.3% | 1.1% | 0.9% |
| Assembly error corrected (Hi-C reads) | 97.9% | 64.7% | 33.2% | 1.2% | 0.9% |
| Assembly duplicate purged | 96.7% | 94.9% | 1.8% | 1.5% | 1.8% |
| Assembly duplicate purged rescued | 97.0% | 95.2% | 1.8% | 1.5% | 1.5% |
| Final Hi-C scaffolded | 98.0% | 96.6% | 1.4% | 0.9% | 1.1% |
| ***Anastrepha ludens*** |  |  |  |  |  |
| Assembly error corrected (long and short reads) | 98.1% | 80.0% | 18.1% | 0.5% | 1.4% |
| Assembly error corrected (Hi-C reads) | 98.0% | 79.9% | 18.1% | 0.6% | 1.4% |
| Assembly duplicate purged | 95.7% | 95.2% | 0.5% | 1.2% | 3.1% |
| Assembly duplicate purged rescued | 96.9% | 96.0% | 0.9% | 1.2% | 1.9% |
| Final Hi-C scaffolded | 97.1% | 96.6% | 0.5% | 1.1% | 1.8% |
| ***Bactrocera dorsalis*** |  |  |  |  |  |
| Assembly error corrected (long and short reads) | 99.2% | 35.4% | 63.8% | 0.4% | 0.4% |
| Assembly error corrected (Hi-C reads) | 99.1% | 35.4% | 63.7% | 0.4% | 0.5% |
| Assembly duplicate purged | 95.2% | 93.5% | 1.7% | 0.8% | 4.0% |
| Assembly duplicate purged rescued | 98.0% | 95.5% | 2.5% | 0.9% | 1.1% |
| Final Hi-C scaffolded | 98.3% | 97.3% | 1.0% | 0.6% | 1.1% |
| ***Bactrocera zonata*** |  |  |  |  |  |
| Assembly error corrected (long and short reads) | 99.3% | 47.3% | 52.0% | 0.2% | 0.5% |
| Assembly error corrected (Hi-C reads) | 99.3% | 47.4% | 51.9% | 0.2% | 0.5% |
| Assembly duplicate purged | 97.8% | 95.4% | 2.4% | 0.7% | 1.5% |
| Assembly duplicate purged rescued | 98.3% | 95.2% | 3.1% | 0.8% | 0.9% |
| Final Hi-C scaffolded | 98.5% | 96.3% | 2.2% | 0.7% | 0.8% |
| ***Zeugodacus cucurbitae*** |  |  |  |  |  |
| Assembly error corrected (long and short reads) | 98.2% | 97.2% | 1.0% | 0.8% | 1.0% |
| Assembly error corrected (Hi-C reads) | 98.3% | 97.3% | 1.0% | 0.8% | 0.9% |
| Assembly duplicate purged | 97.7% | 97.2% | 0.5% | 0.8% | 1.5% |
| Assembly duplicate purged rescued | 98.1% | 97.6% | 0.5% | 0.8% | 1.1% |
| Final Hi-C scaffolded | 98.4% | 98.0% | 0.4% | 0.6% | 1.0% |

**Table 3.** Repeat annotation using Repeat Masker with Dfam database, including Diptera family.

|  | *Anastrepha fraterculus* | *Anastrepha ludens* | *Bactrocera dorsalis* | *Bactrocera zonata* | *Zeugodacus cucurbitae* |
| --- | --- | --- | --- | --- | --- |
| sequences: | 5,502 | 7,879 | 7,027 | 7,749 | 9,971 |
| total length: | 682,452,311 | 721,912,558 | 539,979,902 | 594,029,458 | 374,234,511 |
| GC level: | 37% | 37% | 36% | 36% | 35% |
| bases masked: | 22,626,981<br>(3.2%) | 24,285,344<br>(3.3%) | 17,129,886<br>(3.1%) | 19,389,731<br>(3.1%) | 13,844,259<br>(3.5%) |
|  | number of<br>elements | number of<br>elements | number of<br>elements | number of<br>elements | number of<br>elements |
| Retroelements | 11,922 | 12,534 | 6,950 | 7,529 | 4,606 |
| SINEs: | 508 | 440 | 127 | 157 | 95 |
| Penelope: | 188 | 199 | 136 | 134 | 71 |
| LINEs: | 8,821 | 9,338 | 5,445 | 6,034 | 3,818 |
| L2/CR1/Rex | 4,285 | 4,260 | 1,871 | 2,166 | 725 |
| R2/R4/NeSL | 466 | 490 | 0 | 2 | 0 |
| RTE/Bov-B | 3,707 | 4,153 | 3,377 | 3,664 | 2,972 |
| L1/CIN4 | 346 | 406 | 180 | 169 | 111 |
| LTR elements: | 2,593 | 2,756 | 1,378 | 1,338 | 693 |
| Gypsy/DIRS1 | 2,478 | 2,643 | 1,308 | 1,255 | 642 |
| Retroviral | 110 | 109 | 66 | 72 | 47 |
| DNA transposons | 17,928 | 23,209 | 12,455 | 17,838 | 3,703 |
| hobo-Activator | 1,161 | 1,345 | 1,502 | 1,849 | 394 |
| Tc1-IS630-Pogo | 16,059 | 21,154 | 10,740 | 15,725 | 3,193 |
| MULE-MuDR | 22 | 17 | 6 | 27 | 0 |
| PiggyBac | 346 | 356 | 111 | 109 | 18 |
| Tourist/Harbinger | 1 | 1 | 3 | 3 | 1 |
| Rolling-circles | 1,202 | 1,539 | 822 | 524 | 486 |
| Unclassified: | 2,747 | 3,016 | 2,315 | 2,320 | 830 |
| Small RNA: | 1,756 | 1,544 | 774 | 940 | 635 |
| Satellites: | 229 | 218 | 124 | 139 | 60 |
| Simple repeats: | 295,682 | 311,488 | 241,586 | 262,464 | 226,297 |
| Low complexity: | 51,191 | 53,640 | 33,921 | 39,229 | 31,227 |

**Figure 1.**
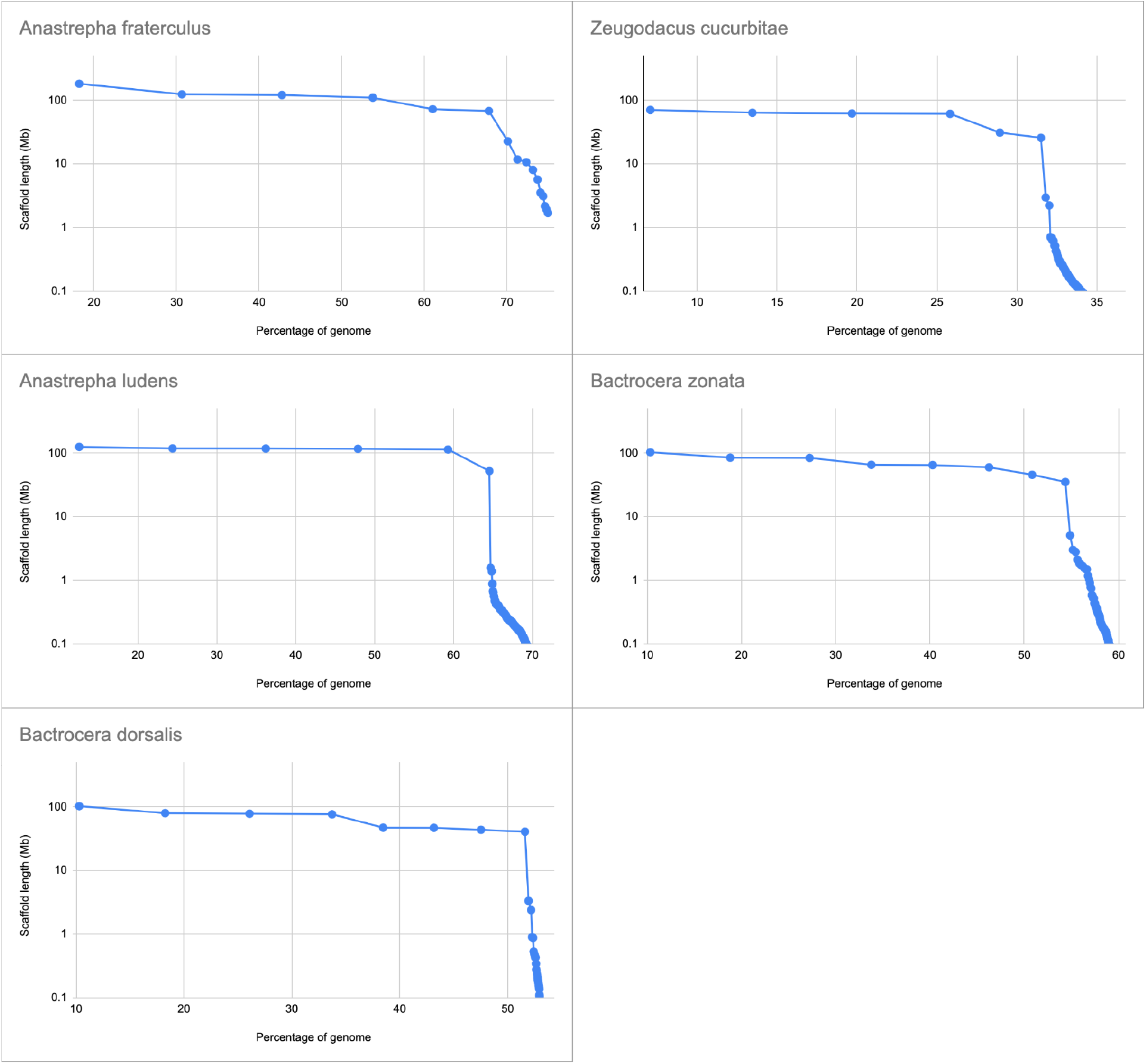
Plot of the percentage of the genome length against the sum of the scaffold sizes adding scaffold from longest to shortest using principal assemblies. To compare the species, we used a common genome size of 1Gb. *Anastrepha fraterculus* (AF), *Anastrepha ludens* (AL), *Bactrocera dorsalis* (BD), *Bactrocera zonata* (BZ) and *Zeugodacus cucurbitae* (ZC). *Anastrepha fraterculus* and *Anastrepha ludens* have their largest contigs sum up to 70% and 65% of the 1 Gb reference size, respectively. *Bactrocera dorsalis* and *Bactrocera zonata* both have their largest contigs sum to 50%-55% of the 1 Gb reference size. *Zeugodacus cucurbitae* assembled to the smallest size at 30% of the 1 Gb reference size.

**Figure 2.**
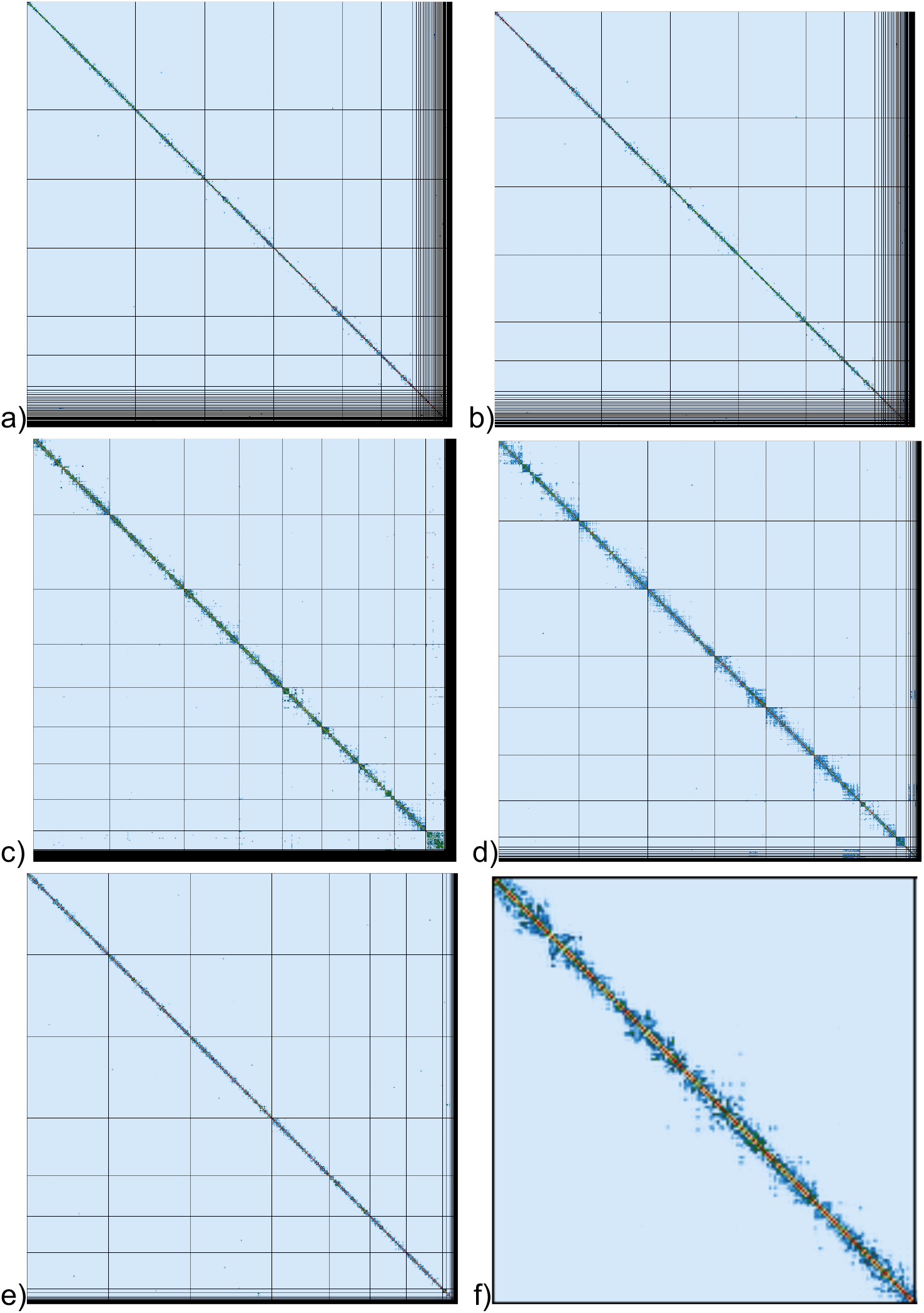
Hi-C maps generated with PretextGraph of the principal assembly for the following species: *Anastrepha fraterculus* (a), *Anastrepha ludens* (b), *Bactrocera dorsalis* (c), *Bactrocera zonata* (d) and *Zeugodacus cucurbitae* (e). In each plot the x-axis and y axis represents the scaffolds of the principal assembly with grid lines separating the scaffolds on both axis. Inside each square, using a heatmap we show the contact map of the interactions in the genome for inter and intra scaffold interactions measured from the HiC read alignments. In panel (f) we show a closeup of the largest primary scaffold for *Zeugodacus cucurbitae* mapped against itself showing the heat map in more detail. As expected, for all the species we mostly see intra-scaffold interactions near the diagonal and no inter-scaffold interactions.

## Discussion

In this work we present assemblies for *Anastrepha fraterculus, Anastrepha ludens, Bactrocera dorsalis, Bactrocera zonata* and *Zeugodacus cucurbitae*. The assemblies were constructed using state-of-the-art technologies, combining both long read sequencing and Hi-C chromatin proximity ligation. Critical to a good initial assembly is the molecular weight and amount of reads used. A large enough contig N50 is important for the Hi-C scaffolding to be successful. A second factor that can impact the Hi-C scaffolding is the presence of duplicated contigs in the assembly as these will cause Hi-C reads to be removed because of duplicate mapping. An optimal duplicate purging step is critical to separate the initial assembly into haploid assemblies and ensure the success of the scaffold sizes into the chromosome scale.

Existing assemblies are deposited in NCBI for *Anastrepha ludens, Bactrocera dorsalis*, and *Zeugodacus cucurbitae*. Although the strains differ and there is merit in each of these assemblies on its own, we compared the BUSCO scores (Table 4) across the assemblies to show that the assemblies presented here are in the same range as in previous studies.

**Table 4.** BUSCO scores across different assembly versions published for *Anastrepha ludens* (AL), *Bactrocera dorsalis* (BD) and *Zeugodacus cucurbitae* (ZC).

| Assembly | BUSCO Complete | BUSCO Duplicated | BUSCO Frag. | BUSCO Missing | # contigs | Total length | N50 | L50 |
| --- | --- | --- | --- | --- | --- | --- | --- | --- |
| *BD (this paper)* | 98.5% | 1.8% | 0.6% | 0.9% | 588 | 537,310,284 | 76,567,395 | 4 |
| BD_GCA_020283865.1 | 92.6% | 0.2% | 0.8% | 6.6% | 6 | 468,671,466 | 90,458,965 | 3 |
| BD_GCA_023373825.1 | 98.1% | 0.2% | 0.8% | 1.1% | 586 | 530,310,706 | 93,297,747 | 3 |
| BD_GCA_029030885.1 | 97.3% | 2.9% | 1.0% | 1.7% | 952 | 524,751,100 | 1,217,930 | 109 |
| BD_GCA_030710565.1 | 95.1% | 1.3% | 0.9% | 4.0% | 1,038 | 566,605,033 | 103,769,376 | 3 |
| *ZC (this paper)* | 98.5% | 0.5% | 0.6% | 0.9% | 5,140 | 374,778,270 | 62,275,020 | 3 |
| ZC_GCA_000806345.2 | 98.2% | 0.2% | 0.7% | 1.1% | 5,545 | 374,633,570 | 1,415,739 | 65 |
| ZC_GCF_028554725.1 | 98.8% | 0.3% | 0.4% | 0.8% | 56 | 439,259,567 | 75,513,634 | 3 |
| *AL (this paper)* | 97.5% | 0.7% | 0.8% | 1.7% | 1,238 | 720,811,084 | 118,249,840 | 3 |
| AL_GCA_029783585.1 | 77.0% | 9.2% | 6.6% | 16.4% | 155,863 | 988,756,442 | 11,016 | 26,222 |
| AL_GCA_028408465.1 | 99.0% | 0.3% | 0.3% | 0.7% | 141 | 820,783,141 | 131,385,309 | 3 |

## Materials and methods

### Sample collection and DNA extraction

Virgin freshly emerged adult males were collected from long-established laboratory colonies of the five fruit fly species at the Insect Pest Control Laboratory of the Joint FAO/IAEA Centre of Nuclear Techniques in Food and Agriculture (Seibersdorf, Austria). The geographic origin and colony generation used for sampling the five species are shown in Supplementary Table S1.

High-Molecular-Weight (HMW) genomic DNA was extracted from the male flies (2-3 flies per extraction) with different kits. The Nanobind tissue kit (Pacific Biosciences) was used for *Anastrepha fraterculus* and *Zeugodacus cucurbitae*. The New England Biolabs (NEB) Monarch DNA extraction kit was used for *Bactrocera dorsalis* and *Bactrocera zonata*. The QIAGEN Genomic tip 20/G kit (Qiagen, Germany) was used for *Anastrepha ludens*. The same extracted DNA was used for long-read sequencing and short-read sequencing.

### Short-read and long-read library construction

The short-read sequencing libraries for all species except *Anastrepha ludens* were constructed using an ultrasonicator (Covaris) for the shearing and the NxSeq AmpFREE Low DNA ligation kit with xGen dual index adapters from Integrated DNA Technologies (IDT). For *Anastrepha ludens* the short read libraries were constructed using TruSeq PCR-free (Illlumina)

The Oxford Nanopore libraries were constructed with unsheared DNA using the Pacific Biosciences short read elimination kit (SRE <25 kb) first and the Oxford Nanopore kit SQK-LSK-109 kit for the library for R9 pore chemistry following the manufacturer’s recommendations.

For the *Anastrepha ludens* PacBio sequencing the sample was purified with AMPure beads (Beckman Coulter, UK) (0.6 volumes) and QC checked for concentration, size, integrity, and purity using Qubit (Qiagen, UK), Fragment Analyser (Agilent Technologies) and Nanodrop (Thermo Fisher) machines. The samples were then processed without shearing using the PacBio Express kit 1 for library construction and an input of 4 µg DNA following the manufacturer’s protocol. The final library was size-selected using the Sage Blue Pippin (Sage Sciences) 0.75% cassette U1 marker in the range of 25–80 kb. The final library size and concentrations were obtained on the Fragment Analyser before being sequenced using the Sequel 1 2.1 chemistry with V4 primers at a loading on plate concentration of 6 pM and 10 h movie times.

### Hi-C proximity ligation library construction

Each proximity ligation library was prepared from several frozen whole flies which were first ground with an SP Bel-Art liquid nitrogen cooled mini mortar (Fisher Scientific). The ground material was then fixed, digested and proximity ligated using the Arima High Coverage HiC+ kit. The final ready-to-sequence library was constructed using a Covaris ultrasonicator for the shearing and the NxSeq AmpFREE Low DNA ligation kit and xGen dual index adapters from Integrated DNA Technologies (IDT).

### Sequencing

The sequencing of the whole genome sequencing libraries was performed on Illumina NovaSeq 6000 S4 (PE150), on Illumina HiSeq 4000 (PE150), and on MGI tech. DNB-G400 (PE150). The Hi-C libraries were sequenced on Illumina NovaSeq 6000 S4 (PE150) and MGI tech. DNB-G400. For MGI sequencing the Illumina-compatible libraries were first circularised for MGI to allow the DNA nanoballs to be generated.

The long read sequencing was performed using both the Pacific Biosciences (PacBio) Sequel and the Oxford Nanopore PromethION P48. See supplementary table S2 for sequencing instruments and sequencing yields. The goat database (goat.genomehubs.org) reports the genome size to be 949Mb for all five species. This genome size was used to calculate the raw read coverage for each type of sequencing.

### Genome assembly and scaffolding

The *Anastrepha ludens* PacBio continuous long reads (CLR) were assembled with the Canu assembler. The other four species (*Anastrepha fraterculus, Bactrocera dorsalis, Bactrocera zonata*, and *Zeugodacus cucurbitae*) sequenced with Oxford Nanopore were assembled with the Flye assembler.

All the assemblies were polished for four rounds using Pilon. The short reads were trimmed with Trimmomatic before being used for polishing. Tigmint was then used with the long reads to further error correct the assemblies. A second correction is performed using the Hi-C data and YaHS by converting the inital_break agp file to a fasta file.

Following the short read, long read, and Hi-C error correction of the assemblies, a haploid version of the assembly is created using Purge_Dups to remove duplicate contigs. This step results in a first version of the principal haploid assembly and an alternate assembly containing the second allele and contig duplicates. To avoid overpurging contigs from the principal haplotype a contig rescue step is performed. The rescue is done by aligning transcripts from the species or closely related species to the principal and alternate assemblies and moving contigs from the alternate to the primary in the case where transcripts only map to the alternate assembly. In the case of *Anastrepha fraterculus* transcripts from *Anastrepha ludens* were used. In the case of *Bactrocera zonata* transcripts from *Bactrocera dorsalis* were used with the addition of the Maleness-on-the-Y (MoY) gene.

At this stage, the main scaffolding is performed on both the principal assembly and on the alternate assembly using chromap for the trimming/mapping and YaHS for the scaffolding. Before scaffolding the alternate assembly, Purge_Dups is used to remove any duplications still present in the alternate assembly. All tool versions used are listed in supplementary table S4. The assembly workflow is illustrated in Figure S1. The version of the tools is listed in supplementary Table S4.

### Additional assembly processing steps for NCBI submission

Before NCBI submission we removed mitochondrial contigs using the following mitochondrial (MT) genome sequences: NC_027725.1 for *Bactrocera zonata*, NC_008748.1 for *Bactrocera dorsalis*, NC_034912.1 for *Anastrepha fraterculus*, MH900082.1 for *Zeugodacus cucurbitae* and MT121222.1 for *Anastrepha ludens*. Scaffolds from the assemblies were aligned to the MT sequences with minimap2 and were removed if more than 80% of the scaffold was aligned. Finally at the end of the workflow the alternative assemblies are missing homozygote scaffolds that were not duplicated in the initial assembly. For NCBI these missing scaffolds were copied back into the alternative assembly to form a more complete alternative assembly.

#### Repeat sequence annotation

The primary assemblies were annotated for repeats using RepeatMasker with the Dfam database (RepeatMasker version 4.1.6, rmblastn version 2.14.1+, Dfam 3.8 with Diptera family).

### Data availability

All sequencing datasets and assemblies have been deposited to National Center for Biotechnology Information (NCBI) in separate Bioprojects for each species.

#### Raw Read Data

The raw sequencing data for the five species have been deposited to NCBI under the sequence read archive (SRA) section. The short read whole genome sequencing (WGS) and Hi-C proximity ligation datasets are provided as untrimmed fastq files. The Oxford Nanopore and PacBio long reads are both provided in fastq format. All the raw read datasets have been submitted under the Bioproject of the corresponding principal assembly (see Table 4).

**Table 4.** Raw sequencing reads are submitted to the NCBI sequence read archive.

| Species | NCBI Bioproject | Dataset type | NCBI SRA accessions |
| --- | --- | --- | --- |
| *Anastrepha fraterculus* | PRJNA1065016 | Long read WGS | SRR29293661, SRR29293662, SRR29293663, SRR29293664 |
|  |  | Short read WGS | SRR29319412, SRR29319413 |
|  |  | Hi-C | SRR29319415, SRR29319414 |
| *Anastrepha ludens* | PRJNA1076526 | Long read WGS | SRR29307005, SRR29307006, SRR29307007 |
|  |  | Short read WGS | SRR29323014, SRR29323013 |
|  |  | Hi-C | SRR29323016, SRR29323015 |
| *Bactrocera dorsalis* | PRJNA1076946 | Long read WGS | SRR29299427 |
|  |  | Short read WGS | SRR29323025, SRR29323026 |
|  |  | Hi-C | SRR29323028, SRR29323027 |
| *Bactrocera zonata* | PRJNA1082643 | Long read WGS | SRR29299502, SRR29299503 |
|  |  | Short read WGS | SRR29319677, SRR29319676 |
|  |  | Hi-C | SRR29319678 |
| *Zeugodacus cucurbitae* | PRJNA1070629 | Long read WGS | SRR29294704, SRR29294702, SRR29294703 |
|  |  | Short read WGS | SRR29322956, SRR29322955 |
|  |  | Hi-C | SRR29322957, SRR29322958 |

### NCBI Assemblies

The contig level principal and alternate assemblies for the five species have been deposited to the NCBI genomes section under separate Bioprojects linked by the same species Biosample (see Table 5). The scaffolded versions are submitted to NCBI.

**Table 5.** Bioprojects and GeneBank assembly IDs in NCBI.

| Species | Biosample | Bioproject (principal) | Bioproject (alternate) | GenBank assembly (principal) | GenBank assembly (alternate) |
| --- | --- | --- | --- | --- | --- |
| *Anastrepha fraterculus* | SAMN39449916 | PRJNA1065016 | PRJNA1065015 | GCA_037575425.2 | GCA_037575645.2 |
| *Anastrepha ludens* | SAMN39944021 | PRJNA1076526 | PRJNA1076525 | GCA_037783455.2 | GCA_037783485.2 |
| *Bactrocera dorsalis* | SAMN39957830 | PRJNA1076946 | PRJNA1076945 | GCA_037783525.2 | GCA_037783465.2 |
| *Bactrocera zonata* | SAMN40214030 | PRJNA1082643 | PRJNA1082642 | GCA_037783105.2 | GCA_037783125.2 |
| *Zeugodacus cucurbitae* | SAMN39655635 | PRJNA1070629 | PRJNA1070628 | GCA_037783285.2 | GCA_037783305.2 |

## Acknowledgments

This study benefited from discussions at meetings for the Coordinated Research Project D44003, “Generic approach for the development of genetic sexing strains for SIT applications”, funded by the International Atomic Energy Agency (IAEA). The authors also wish to thank Elena Isabel Cancio Martinez and Gülizar Pillwax for insect rearing.

## Supplementary Information

**Supplementary Figure S1.**
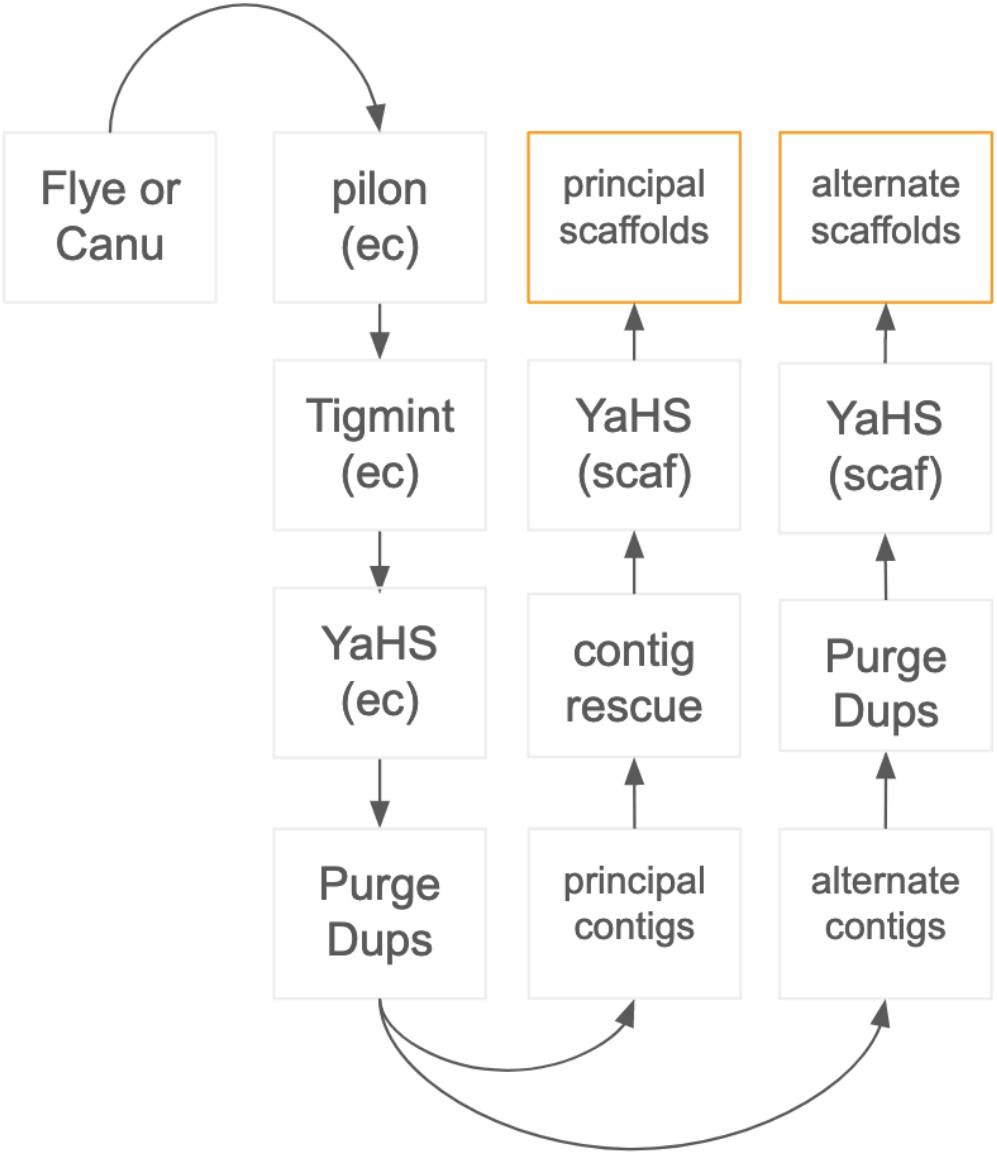
Assembly workflow

**Supplementary Table S1.** Description of fly strains and generations used for the DNA extraction.

| Species | Generation | Geographic origin (strain) |
| --- | --- | --- |
| *Anastrepha fraterculus* | G63 | Brazil (Vacaria) |
| *Anastrepha ludens* | G117 | Mexico |
| *Bactrocera dorsalis* | G113 | Thailand (Saraburi) |
| *Bactrocera zonata* | G17 | Israel |
| *Zeugodacus cucurbitae* | G127 | Mauritius (Sookar 5/a) |

**Supplementary Table S2.** Sequencing yields and genome coverages.

| Species | DataSet Type | Sequencer | Total reads | Total bases | Taxon ID | Genome Size in Mb | Genome fold Coverage |
| --- | --- | --- | --- | --- | --- | --- | --- |
| *Anastrepha fraterculus* | Long read WGS | PrometION R9 | 18,054,709 | 38,607,090,382 | 95504 | 949 | 40.7 |
|  | Short read WGS | NovaSeq 6000, S4 | 55,815,810 | 16,856,374,620 |  |  | 17.8 |
|  | Hi-C | NovaSeq 6000, S4 | 261,079,890 | 78,323,967,000 |  |  | 82.5 |
| *Anastrepha ludens* | Long read WGS | PacBio, Sequel | 2,354,154 | 29,476,521,382 | 28586 | 949 | 31.1 |
|  | Short read WGS | HiSeq 4000 | 62713989 | 18486278987 |  |  | 19.5 |
|  | Hi-C | NovaSeq 6000, S4 | 172,962,558 | 51,888,767,400 |  |  | 54.7 |
| *Bactrocera dorsalis* | Long read WGS | PrometION R9 | 4,135,925 | 74,158,935,467 | 27457 | 949 | 78.1 |
|  | Short read WGS | HiSeq 4000 | 49,171,609 | 14,511,740,112 |  |  | 15.3 |
|  | Hi-C | NovaSeq 6000, S4 | 288,365,720 | 86,509,716,000 |  |  | 91.2 |
| *Bactrocera zonata* | Long read WGS | PrometION R9 | 2,172,892 | 38,457,368,805 | 137042 | 949 | 40.5 |
|  | Short read WGS | MGI DNB-G400 | 110,367,330 | 33,110,199,000 |  |  | 34.9 |
|  | Hi-C | MGI DNB-G400 | 154,157,696 | 46,247,308,800 |  |  | 48.7 |
| *Zeugodacus cucurbitae* | Long read WGS | PrometION R9 | 21,084,151 | 87,690,269,635 | 28588 | 949 | 92.4 |
|  | Short read WGS | NovaSeq 6000, S4 | 60,261,930 | 18,199,102,860 |  |  | 19.2 |
|  | Hi-C | NovaSeq 6000, S4 | 154,157,696 | 46,247,308,800 |  |  | 48.7 |

**Supplementary Table S3.** Software tool versions.

| Tool | Version | Software Link |
| --- | --- | --- |
| Canu | 1.8 | https://github.com/marbl/canu |
| Flye | 2.9-b1774 | https://github.com/fenderglass/Flye |
| pilon | 1.23 | https://github.com/broadinstitute/pilon |
| Trimmomatic | 0.36 | http://www.usadellab.org/cms/?page=trimmomatic |
| Tigmint | 1.2.10 | https://github.com/bcgsc/tigmint |
| Purge_Dups | 1.2.5 | https://github.com/dfguan/purge_dups |
| chromap | 0.2.5-r473 | https://github.com/haowenz/chromap |
| YaHS | 1.2 | https://github.com/c-zhou/yahs |
| fcs-adaptor | 0.5.0 | https://github.com/ncbi/fcs |
| minimap2 | 2.12 | https://github.com/lh3/minimap2 |

## Literature Cited

Ahn JJ, Choi KS, Huang Y-B. 2022. Thermal effects on the development of Zeugodacus cucurbitae (Coquillett) (Diptera: Tephritidae) and model validation. Phytoparasitica. 50:601–616. doi: 10.1007/s12600-022-00985-5.

Augustinos AA et al. 2017. Ceratitis capitata genetic sexing strains: laboratory evaluation of strains from mass-rearing facilities worldwide. Entomol Exp Appl. 164:305–317. doi: 10.1111/eea.12612.

Bai X et al. 2019. CRISPR/Cas9-mediated knockout of the eye pigmentation gene white leads to alterations in colour of head spots in the oriental fruit fly, Bactrocera dorsalis. Insect Molecular Biology. 28:837–849. doi: 10.1111/imb.12592.

CABI. 2019. Invasive Species Compendium. Datasheet Anastrepha ludens (Mexican fruit fly) 19/11/1919. Wallingford, UK: CAB International. Available online: https://www.cabi.org/isc/datasheet/5654.

Caceres C. 2002. Mass Rearing of Temperature Sensitive Genetic Sexing Strains in the Mediterranean Fruit Fly (Ceratitis Capitata). Genetica. 116:107–116. doi: 10.1023/A:1020967810703.

De Meyer M et al. 2015. A review of the current knowledge on Zeugodacus cucurbitae (Coquillett) (Diptera, Tephritidae) in Africa, with a list of species included in Zeugodacus. ZK. 540:539–557. doi: 10.3897/zookeys.540.9672.

Delrio G, Cocco A. 2012. THE PEACH FRUIT FLY, BACTROCERA ZONATA: A MAJOR THREAT FOR MEDITERRANEAN FRUIT CROPS? Acta Hortic. 557–566. doi: 10.17660/ActaHortic.2012.940.80.

Dhillon MK, Singh R, Naresh JS, Sharma HC. 2005. The melon fruit fly, Bactrocera cucurbitae: A review of its biology and management. Journal of Insect Science. 5. doi: 10.1093/jis/5.1.40.

Dyck VA, Hendrichs J, Robinson AS, eds. 2021. Sterile Insect Technique: Principles and Practice in Area-Wide Integrated Pest Management. 2nd ed. CRC Press doi: 10.1201/9781003035572.

El-Mahdy SM, Amin A-RH, Emam AK, Hashem A-FG. 2009. BIOLOGICAL STUDIES OF THE PEACH FRUIT FLY, BACTROCERA ZONATA (SAUNDERS) (TEPHRITIDAE: DIPTERA) ON THREE ARTIFICIAL DIETS AT THREE CONSTANT TEMPERATURES. Egyptian Journal of Agricultural Research. 87:1025–1040. doi: 10.21608/ejar.2009.198817.

Enkerlin WR. 2005. Impact of Fruit Fly Control Programmes Using the Sterile Insect Technique. In: Sterile Insect Technique. Dyck, VA, Hendrichs, J, & Robinson, AS, editors. Springer-Verlag: Berlin/Heidelberg pp. 651–676. doi: 10.1007/1-4020-4051-2_25.

EPPO. 2024. EPPO Global Database.

FAO/IAEA. 2019. Report of the First Research Coordination Meeting on the “Generic approach for the development of genetic sexing strains for sterile insect technique applications.”. https://www.iaea.org/sites/default/files/20/11/d44003-rcm1report_20200304_0.pdf.

FAO/IAEA. 2021. Report of the Second Research Coordination Meeting on the “Generic approach for the development of genetic sexing strains for sterile insect technique applications.”. d44003-crp_rcm2-report.pdf (iaea.org).

Franz G, Bourtzis K, Cáceres C. 2021. Practical and Operational Genetic Sexing Systems Based on Classical Genetic Approaches in Fruit Flies, an Example for other Species Amenable to Large-Scale Rearing for the Sterile Insect Technique. In: Sterile Insect Technique Principles and Practice in Area-Wide Integrated Pest Management. https://www.taylorfrancis.com/chapters/oa-edit/10.1201/9781003035572-17/practical-operational-genetic-sexing-systems-based-classical-genetic-approaches-fruit-flies-example-species-amenable-large-scale-rearing-sterile-insect-technique-franz-bourtzis-c%C3%A1ceres?context=ubx&refId=318df6bb-c6c9-4e99-94a6-0dabb26be636.

Hendrichs J, Franz G, Rendon P. 1995. Increased effectiveness and applicability of the sterile insect technique through male-only releases for control of Mediterranean fruit flies during fruiting seasons. J Applied Entomology. 119:371–377. doi: 10.1111/j.1439-0418.1995.tb01303.x.

Hendrichs J, Vera T, De Meyer M, Clarke A. 2015. Resolving cryptic species complexes of major tephritid pests. ZK. 540:5–39. doi: 10.3897/zookeys.540.9656.

Hernández E et al. 2017. The Effects of a Modified Hot Water Treatment on Anastrepha ludens (Diptera: Tephritidae)-Infested Mango. J Econ Entomol. tow245. doi: 10.1093/jee/tow245.

Jaffar S, Rizvi SAH, Lu Y. 2023. Understanding the Invasion, Ecological Adaptations, and Management Strategies of Bactrocera dorsalis in China: A Review. Horticulturae. 9:1004. doi: 10.3390/horticulturae9091004.

Khan RA, Naveed M. 2017. Occurrence and Seasonal Abundance of Fruit Fly, Bactrocera zonata Saunders (Diptera: Tephritidae) in Relation to Meteorological Factors. PJZ. 49:999– 1003. doi: 10.17582/journal.pjz/2017.49.3.999.1003.

Klassen W, Curtis CF, Hendrichs J. 2021. History of the Sterile Insect Technique. In: Sterile Insect Technique Principles and Practice in Area-Wide Integrated Pest Management. CRC Press.

Komal J et al. 2023. Unveiling the Genetic Symphony: Harnessing CRISPR-Cas Genome Editing for Effective Insect Pest Management. Plants. 12:3961. doi: 10.3390/plants12233961.

Ovruski S, Schliserman P, Aluja M. 2003. Native and Introduced Host Plants of Anastrepha fraterculus and Ceratitis capitata (Diptera: Tephritidae) in Northwestern Argentina. Journal of Economic Entomology. 96:1108–1118. doi: 10.1093/jee/96.4.1108.

Papadopoulos NT, De Meyer M, Terblanche JS, Kriticos DJ. 2024. Fruit Flies: Challenges and Opportunities to Stem the Tide of Global Invasions. Annu. Rev. Entomol. 69:355–373. doi: 10.1146/annurev-ento-022723-103200.

Paulo DF et al. 2022. A Unified Protocol for CRISPR/Cas9-Mediated Gene Knockout in Tephritid Fruit Flies Led to the Recreation of White Eye and White Puparium Phenotypes in the Melon Fly Tamborindeguy, C, editor. Journal of Economic Entomology. 115:2110–2115. doi: 10.1093/jee/toac166.

Qin Y, Wang C, Zhao Z, Pan X, Li Z. 2019. Climate change impacts on the global potential geographical distribution of the agricultural invasive pest, Bactrocera dorsalis (Hendel) (Diptera: Tephritidae). Climatic Change. 155:145–156. doi: 10.1007/s10584-019-02460-3.

Rendón P, McInnis D, Lance D, Stewart J. 2004. Medfly (Diptera:Tephritidae) Genetic Sexing: Large-Scale Field Comparison of Males-Only and Bisexual Sterile Fly Releases in Guatemala. ec. 97:1547–1553. doi: 10.1603/0022-0493-97.5.1547.

Segura DF et al. 2006. Relative Abundance of *Ceratitis capitata* and *Anastrepha fraterculus* (Diptera: Tephritidae) in Diverse Host Species and Localities of Argentina. an. 99:70–83. doi: 10.1603/0013-8746(2006)099[0070:RAOCCA]2.0.CO;2.

Sollazzo G et al. 2024. Deep orange gene editing triggers temperature-sensitive lethal phenotypes in Ceratitis capitata. BMC Biotechnol. 24:7. doi: 10.1186/s12896-024-00832-x.

Ullah F et al. 2023. Estimation of the potential geographical distribution of invasive peach fruit fly under climate change by integrated ecological niche models. CABI Agric Biosci. 4:46. doi: 10.1186/s43170-023-00187-x.

Vargas R, Piñero J, Leblanc L. 2015. An Overview of Pest Species of Bactrocera Fruit Flies (Diptera: Tephritidae) and the Integration of Biopesticides with Other Biological Approaches for Their Management with a Focus on the Pacific Region. Insects. 6:297–318. doi: 10.3390/insects6020297.

Ward CM et al. 2021. White pupae phenotype of tephritids is caused by parallel mutations of a MFS transporter. Nat Commun. 12:491. doi: 10.1038/s41467-020-20680-5.

White IM, Elsonharris MM. 1994. Fruit Flies of Economic Significance: Their Identification and Bionomics. Environmental Entomology. 22:1408–1408. doi: 10.1093/ee/22.6.1408a.

Zapata S. 2022. Economic Implications of the Mexican Fruit Fly Infestation in Texas. JOE. 60. doi: 10.34068/joe.60.02.03.

Zhang Y et al. 2023. Genomes of the cosmopolitan fruit pest Bactrocera dorsalis (Diptera: Tephritidae) reveal its global invasion history and thermal adaptation. Journal of Advanced Research. 53:61–74. doi: 10.1016/j.jare.2022.12.012.

Zhao Z, Carey JR, Li Z. 2024. The Global Epidemic of Bactrocera Pests: Mixed-Species Invasions and Risk Assessment. Annu. Rev. Entomol. 69:219–237. doi: 10.1146/annurev-ento-012723-102658.

Zingore KM et al. 2020. Global risk of invasion by Bactrocera zonata: Implications on horticultural crop production under changing climatic conditions Yue, B-S, editor. PLoS ONE. 15:e0243047. doi: 10.1371/journal.pone.0243047.

Angela Meccariello et al. Maleness-on-the-Y (MoY) orchestrates male sex determination in major agricultural fruit fly pests. Science 27 Sep 2019 Vol 365, Issue 6460 pp. 1457–1460 DOI: 10.1126/science.aax1318

Rhie, A., McCarthy, S.A., Fedrigo, O. et al. Towards complete and error-free genome assemblies of all vertebrate species. Nature 592, 737–746 (2021). 10.1038/s41586-021-03451-0

Felipe A. Simao et al. BUSCO: assessing genome assembly and annotation completeness with single-copy orthologs. Bioinformatics, 31(19), 2015, 3210–3212 doi: 10.1093/bioinformatics/btv351

